# Reevaluating Maturity and Spawning Potential of Atlantic Bluefin Tuna in the Slope Sea

**DOI:** 10.64898/2026.08.04.742198

**Authors:** Chi Hin Lam, Gilad Heinisch, Aldo Corriero, Molly Lutcavage

## Abstract

The management of Atlantic bluefin tuna (*Thunnus thynnus*) is currently defined by a long-standing life-history paradox: a stark divergence in maturation schedules between the Eastern and Western stocks. While the Mediterranean contingent matures at approximately 104 cm straight fork length (SFL; age 3-5), the western stock has long been characterized as a late-maturing, at ∼190 cm SFL (age 8+), and as late as age 15.8, based on size-at-catch data in the presumed exclusive spawning areas of the Gulf of Mexico. This discrepancy defies maturity states confirmed by endocrine hormones as well as established life-history principles that link metabolic energetics to reproductive biology. To resolve this critical issue in stock assessment, we present a sensitivity analysis based on the first histological evidence of spawning- capable individuals from 42 bluefin females (97-244 cm curved fork length or ∼95- 236 cm SFL) sampled in June 2025 within the Slope Sea, Northwest Atlantic. This historically overlooked, temperate spawning ground must be recognized and past estimates of stock productivity reevaluated.

Our data provide a conservative estimate of length-at-50% maturity, L_50_ for females at 123.3 cm SFL, effectively reconciling the reproductive parameters of the two stocks based on histology. Our findings align established bluefin tuna metabolic and evolutionary symmetries in life history, irrespective of management boundaries. Incorporating this previously unrecognized spawning group (i.e., smaller, younger fish) calls for a substantial upward revision of Western spawning stock biomass (SSB). This biomass represents an intrinsic biological buffer that could contribute to the Atlantic bluefin’s adaptation and resilience. Appreciating demographic diversity is critical to accurately diagnosing stock vulnerability, ensuring that management frameworks protect long-term population stability in the face of climate-driven oceanographic shifts and ongoing exploitation pressure.

## Introduction

Fisheries management of the Atlantic bluefin tuna (*Thunnus thynnus*, hereafter ABFT) by the International Commission for the Conservation of Atlantic Tunas (ICCAT) is tethered to a life-history paradigm established decades ago. This paradigm stipulates late maturity (age-8+, >190 cm straight fork length or SFL) and exclusive spawning within the Gulf of Mexico and around the Florida Straits [Rivas 1954; Baglin 1982; Farley and Ohshimo 2019], tenets routinely invoked in past and current stock assessments. However, this ‘late-maturing’ model was based on biological assumptions at odds with multiple sources of historic (e.g., Wilson and Bartlett 1967) and recent empirical evidence [Goldstein et al. 2007, Heinisch et al. 2014, and more reviewed in Corriero et al. 2020], and life-history modeling [Chapman et al. 2011]. The persistent presence of larvae in the Slope Sea, a temperate region between the Gulf Stream and the U.S. continental shelf, is unambiguous evidence of reproductive activities occurring outside the Gulf of Mexico [Richardson et al., 2016; 2026]. Together, these datasets challenge the long-standing “late-maturity” paradigm.

While initial Slope Sea rediscoveries were rejected by some as inconsequential [Walter et al. 2016] or met with unfounded skepticism [Safina 2016], the pivotal question has since shifted from whether spawning occurs in the Slope Sea to the stock identity of these smaller adults and their demographic relationship to other spawning grounds. To date, persistent ambiguity regarding the origin and maturity of Slope Sea adults has stalled the integration of this spawning habitat into the fisheries management process. Previous interpretations have characterized the region as a transient ‘outpost’ or mixing zone for migratory contingents of both eastern and western origin [Walter et al. 2016]. By framing the Slope Sea as a simple dispersal corridor, and ignoring established sexual maturity for smaller fish [Heinisch et al. 2014], historical stock assessments have overlooked the region’s potential as a vital spawning habitat, despite early fishery observations [Wilson and Bartlett 1967; Mather et al. 1995] and biotelemetry-based predictions of adult occupancy [Lutcavage et al. 1999; Galuardi et al. 2010; Galuardi and Lutcavage 2012].

This study reframes the depiction of ABFT productivity by evaluating the role of the Slope Sea as a recurring spawning ground. We integrate maturity modeling of 42 female tuna sampled in the Slope Sea during June 2025 (Lutcavage et al. *in prep*) with sensitivity analyses of maturity-at-length parameters and considerations of stock identity. Our results reveal that the Slope Sea supports a productive, spawning contingent that is omitted in ICCAT management frameworks. We demonstrate that recognizing this habitat is essential for realistic understanding of ABFT, regardless of mixed-stock proportions or management boundaries.

## Methods

### Field Sampling

Biological sampling of adult ABFT was conducted around the Northeast Canyons (39.75**°**–39.93**°**N, 69.23**°**–71.43**°**W) in the Slope Sea from 16-27 June 2025. Gonads (n = 90; 42 females, 48 males) were collected aboard the F/V *Eagle Eye II* during multi-species commercial longline operations [Lutcavage et al., *in prep*]. To ensure regulatory compliance, our sampling was limited to the size-class allocations and landing restrictions of U.S. Highly Migratory Species Exempted Fishing Permit HMS- EFP-25-24 (EFP). Consequently, predefined permit parameters established the operational limits of our sampling. Specifically, retention was capped at a quota of 60 individuals (combined sexes) within the 150-185 cm curved fork length (CFL) size class. Once EFP allocations were reached, irrespective of continued availability, ABFT were either tagged and released (n=24), processed as mandatory regulatory discards, or retained as catch allowed under the vessel’s Individual Bluefin Quota during active commercial operations.

### Histological Maturity Classification

Biological sampling and histological protocols are detailed in Lutcavage et al. (*in prep*). Following Corriero et al. (2003), females were classified as mature based on the presence of advanced oocyte developmental stages, including vitellogenesis, migratory nucleus, or post-ovulatory follicles (POFs). “Active spawning” was specifically identified by the presence of hydrated oocytes or POFs, indicating spawning within 24-48 hours of capture.

### Comparative Maturity Analysis

The 2025 cruise observations were contrasted against two established maturity paradigms:

1. A western Atlantic baseline based on an operational, knife-edge threshold at L_50_ of 190.0 cm SFL, utilized in ICCAT assessments [Anon. 2009; 2017; 2021; 2022]. This assumption of late maturity was based on reproductive sampling conducted by Baglin (1982) between 1974 and 1978 [Farley and Ohshimo 2019] and size-at-catch based evaluations for fish caught in the Gulf of Mexico [Nemerson et al. 2000; Diaz and Turner 2007; Diaz 2011]. The Baglin (1982) historic dataset included small-to-medium fish in the Mid-Atlantic Bight (June-August) and large adults (’giants’) in the Gulf of Mexico (March-June), Bahamas (April-June), and the U.S. Northeast (July-October).
2. The Mediterranean-derived model [Corriero et al. 2005], a maturity ogive representing the Eastern Atlantic and Mediterranean life history, where sexual maturation is reached by age 3-5. This paradigm is characterized by an L_50_ of 103.6 cm SFL, reflecting earlier sexual maturation than the western Atlantic model. Regarded as a benchmark in reproductive profiling, this model’s robustness is based on a comprehensive biological dataset (n = 501 females) with definitive histological staging.

Length (L) measurements measured as curved fork length for Slope Sea ABFT were standardized to straight fork length (SFL). Curved fork length (CFL) was converted to SFL using conversion factors listed in ICCAT manual [ICCAT 2006-2016]. Weight-at-length (W) was calculated as

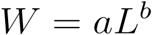

where a = 1.77054 x 10^5^ and b = 3.001252 [ICCAT 2006-2016].

### Spawning Stock Biomass and Maturity Modeling

To quantify the impact of maturity assumptions on stock perception, we modeled total spawning stock biomass (SSB) as an integrated function of length from 60-300 cm SFL:

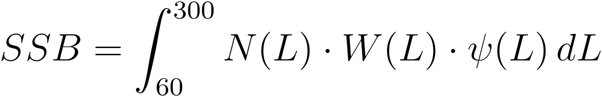

Numerical abundance (N) followed an exponential length-based decay:

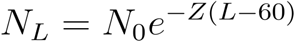

where Z = 0.02 cm^-1^ served as a length-based mortality proxy [Charnov 1993] consistent with the age-specific natural mortality (M ∼ 0.14) adopted by the ICCAT SCRS [Anon. 2009; 2021; 2022]. The probability of maturation at length (ψ_L_) followed a logistic fit with a fixed slope (β= 0.1739) derived from Corriero et al. (2005):

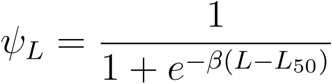

Sensitivity analysis was performed across a range of L_50_ values (100.3 cm to 250 cm), with 123.3 cm serving as a research-based threshold to quantify reproductive potential below the historical 190-cm management cutoff.

## Results

Biological sampling of 42 female ABFT in the Slope Sea yielded a size distribution from 95 to 236 cm SFL (Fig. 1). The observed length frequencies of our reproductive sampling reflect both the localized availability of the catch and the strict regulatory constraints of our Exempted Fishing Permit. The resulting histological evaluation of ovaries demonstrated that 93% (n = 39) of all females sampled were sexually mature, including every individual exceeding 120 cm SFL (Lutcavage et al. *in prep*). All sampled males were likewise determined to be sexually mature (Lutcavage et al. *in prep*); however, for stock assessment purposes, subsequent analyses focus primarily on female reproductive parameters.

**Figure 1.**
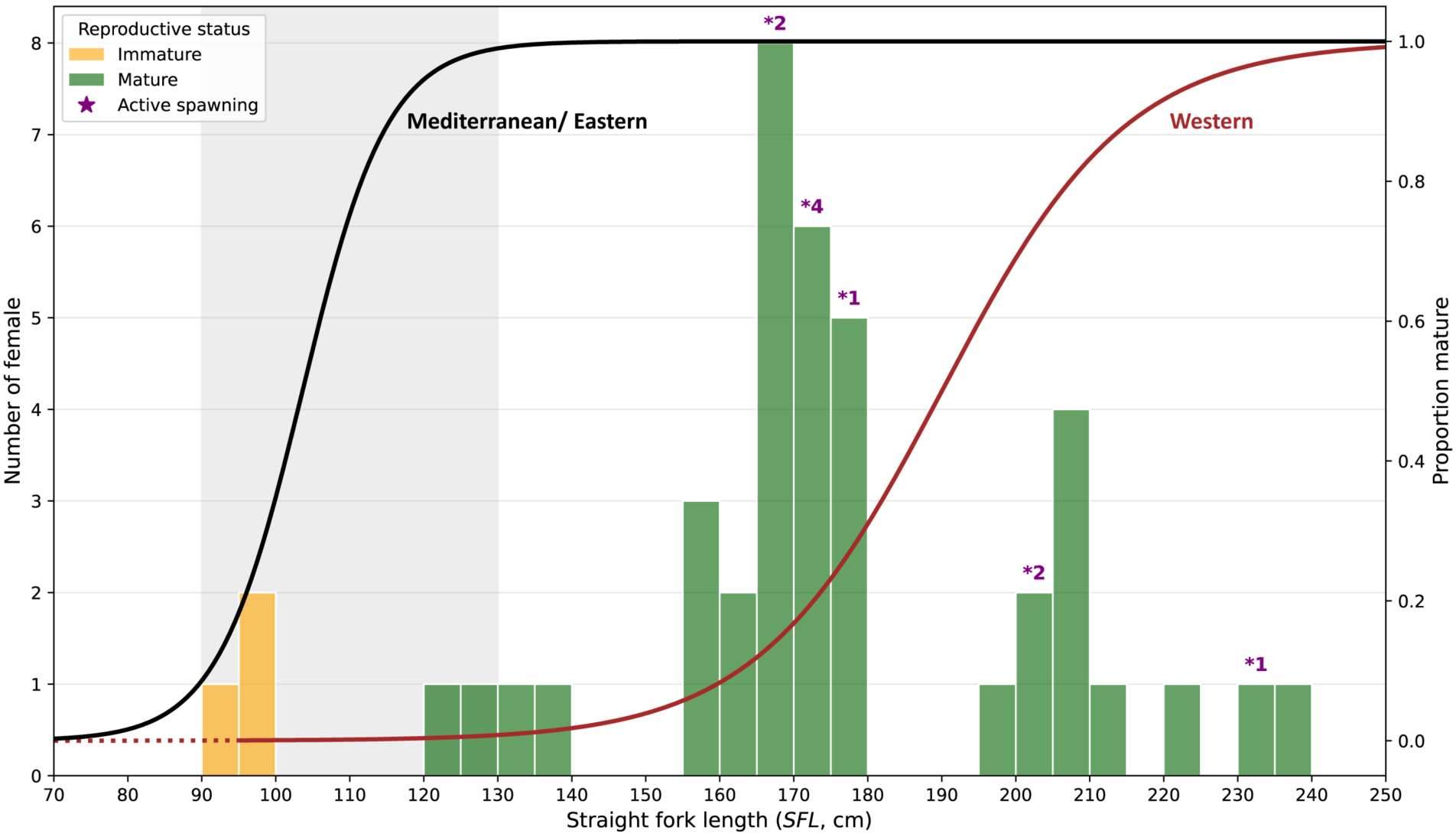
Comparative Maturity-at-Length Analysis for Atlantic Bluefin Tuna, Western Atlantic. Histological observations of 42 females from the 2025 sampling cruise (stacked histogram) align with the Mediterranean-derived maturity model (Corriero et al. 2005; solid black line, length-at-50% maturity L_50_ = 103.6 cm straight fork length, SFL). Individual reproductive status was classified as immature (orange) or mature (dark green). Counts of actively spawning females (168-231 cm SFL, n = 10), shown above the corresponding size classes. These data are contrasted with the western Atlantic baseline (brown line, L_50_ = 190.0 cm SFL), based on Baglin (1982). A dotted line between 70 and 95 cm SFL denotes extrapolation below Baglin’s minimum length (95 cm SFL) from 81 females sampled in the Mid-Atlantic Bight. The light grey shaded region (90–130 cm SFL) highlights a critical sampling zone missing in historic data. Note a slight offset is added to the secondary y-axis, ‘Proportion mature’.

Females between 130 and 190 cm SFL are currently classified as immature within past ICCAT assessments [e.g., Anon. 2009; 2017; 2021; 2022]. Contrary to this assumption, we identified active spawning in 10 females (24% of females sampled; 168–231 cm SFL). Notably, these spawning individuals were captured across five distinct longline set locations spanning the 11-day fishing trip. This persistent presence of spawning-capable ABFT of both sexes across the vast majority of our sampling effort (9 out of 10 longline sets; 1 set was truncated to two hours due to operational challenges) establishes a direct, empirical biological link between adult reproductive activity and regional larval collections [Richardson et al. 2016; 2026].

### Spawning Stock Biomass Sensitivity to Maturity Thresholds

Simulated spawning stock biomass (SSB) exhibited high sensitivity to the selection of the L_50_ maturity threshold (Fig. 2A). Transitioning from the late-maturation baseline (L_50_ = 190 cm) to our empirically derived threshold (L_50_ = 123.3 cm) resulted in a 90% SSB increase. This shift demonstrates that the reproductive capacity of the stock is nearly double what is currently estimated under late-maturity assumptions.

**Figure 2.**
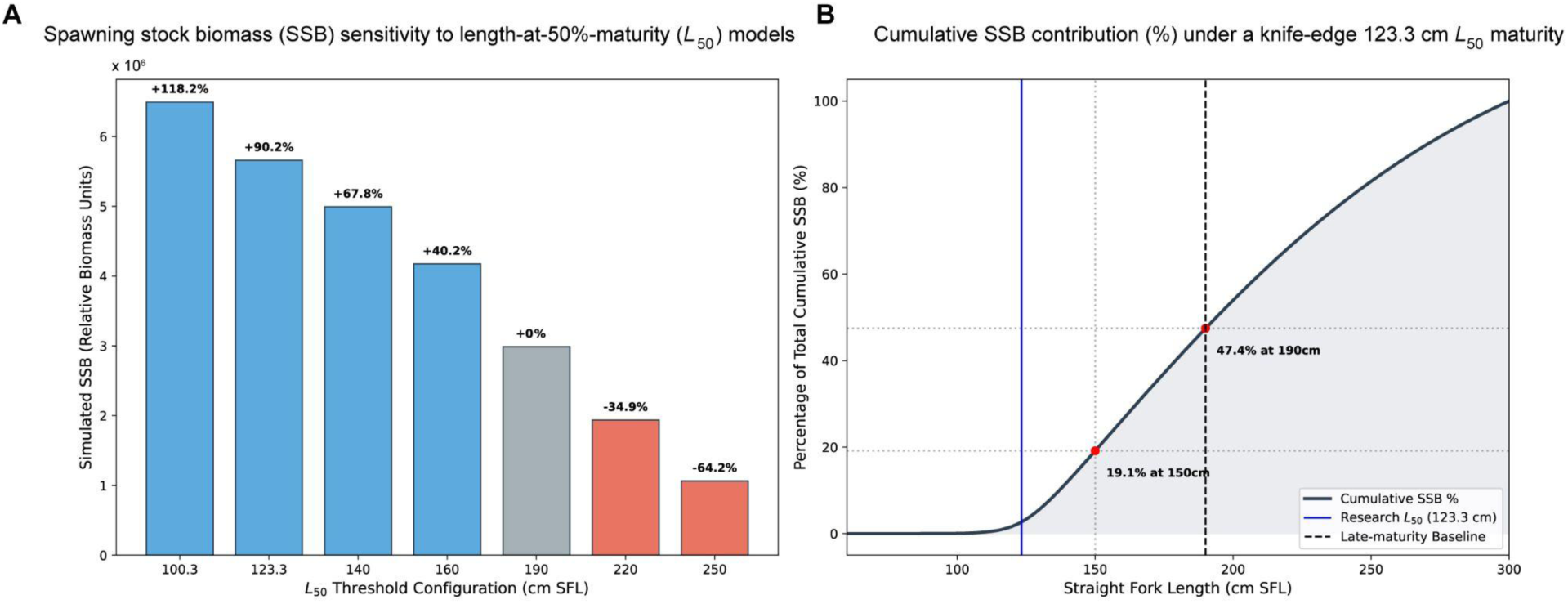
Spawning stock biomass (SSB) sensitivity and distribution. (A) Sensitivity of total simulated SSB to variations in the length-at-50% maturity (L_50_; straight fork length, SFL, cm), calculated using Corriero et al. (2005) maturity ogive slope (β=0.1739). (B) Cumulative distribution of SSB under the 123.3 cm L_50_ research-based threshold. The shaded region highlights that 47% of spawning biomass is composed of younger, mid-sized cohorts (<190 cm) omitted from management frameworks.

The cumulative SSB distribution highlights the substantial contribution of mid-sized fish to the total reproductive potential (Fig. 2B). Under our biological findings, i.e., research-based threshold model, fish between 123.3 cm and 190 cm SFL account for 47% of the total SSB. The steepness of the cumulative curve across this specific range indicates that the population’s total reproductive potential is heavily supported by these younger adults, where the higher numerical abundance of individuals effectively compensates for lower individual batch fecundity relative to larger, older cohorts [Medina et al. 2002; 2007; 2016; Aranda et al. 2013; Knapp et al. 2014; Addis et al. 2016].

## Discussion

Our documentation of spawning-capable females as small as 120 cm SFL calls for a fundamental re-evaluation of the ABFT western stock reproductive schedule. This revised schedule is independently corroborated by Stewart et al. (2024), who identified a growth-based maturation transition at approximately age 4 (∼116 cm SFL) common to both Atlantic stocks. Ultimately, this alignment reconciles the stock’s modeled reproductive potential with its documented larval distribution [Richardson et al. 2016; 2026] and occupancy by smaller adults in the Slope Sea [Galuardi and Lutcavage 2012].

By establishing this unified maturity schedule, the convergence of physiological evidence [this study; Goldstein et al. 2007; Heinisch et al. 2014; Knapp et al. 2014] and life history modeling [Chapman et al. 2011] provides a rigorous rebuttal to the flawed practice of using size-at-catch data to determine a physiological process, namely, sexual maturity. Specifically, these data challenge conclusions of delayed maturity derived from biotelemetry tracks [Aalto et al. 2023] or length-frequency profiles restricted to the Gulf of Mexico [Nemerson et al. 2000; Diaz and Turner 2007; Diaz 2011]. Our histological findings provide the definitive biological validation for the otolith results of Stewart et al. (2024), demonstrating that the Western stock’s functional spawning contingent extends far below the current paradigm of a 190-cm threshold.

### The Biological Basis of Population Resilience

Our results provide a mechanistic explanation for the demographic resilience of the Western stock. Traditionally, stock stability was attributed exclusively to highly fecund, older cohorts exceeding age 8 (≥190 cm SFL). However, our findings reveal that nearly half of the total reproductive potential is supported by younger, mid-sized individuals. This distributed spawning potential provides a crucial biological buffer, maintaining population productivity even during periods of truncated age structures, or reduced biomass in the oldest age classes.

To understand the demographic impact of these younger females, it is critical to distinguish between the maturity slope parameter (β), which governs the physiological transition of cohorts into the spawning population, and stock-recruitment steepness (h), which characterizes the demographic response to population depletion [Mangel et al. 2010]. While the maturity schedule defines the absolute magnitude and size composition of spawning stock biomass, it is distinct from the density-dependent feedback mechanisms that govern subsequent recruitment.

The rapid accumulation of SSB within the 123.3-190.0 cm SFL range indicates that these intermediate cohorts are the primary drivers of stock productivity. Although a 250-cm “giant” may have exponentially higher absolute fecundity, the aggregate reproductive output of the far more numerous mid-sized fish (130-190 cm SFL) provides a more stable, diversified reproductive portfolio. Consequently, the Western stock possesses a higher intrinsic rate of population increase (r) than previously recognized. This confers an enhanced capacity to withstand oceanographic fluctuations and anthropogenic perturbations- e.g., climate change, disease, prey dynamics, and overfishing [Lowerre-Barbieri et al. 2017].

### The Portfolio Effect and Spatiotemporal Bet-Hedging

Re-positioning of the Slope Sea as a recurring, productive spawning ground advances our understanding of *T. thynnus*. These findings shift the paradigm from a vulnerable, late-maturing Western stock [Block et al. 2026] toward a new framework defined by size-structured migration and spawning-site heterogeneity. This update represents a classic ecological “portfolio effect” [Schindler et al. 2010; Richardson et al. 2016], wherein total reproductive effort is distributed across distinct oceanographic regimes, ranging from oligotrophic, subtropical waters of the Gulf of Mexico and Caribbean area to the nutrient-rich, temperate frontal systems of the Slope Sea. Analogously, Pacific bluefin tuna (*Thunnus orientalis*) demonstrate similar demographic and energetic profiles [Chen et al. 2006; Itoh, 2006], and achieve the decoupling of recruitment success from the environmental conditions of a single geographic location. In this respect, temperate bluefin tuna species achieve a robust level of spatiotemporal bet-hedging. For a long-lived marine vertebrate, this reproductive strategy likely dampens destabilizing impacts of environmental stochasticity and localized anomalies on annual recruitment.

### Genomic Validation of Stock Identity

A central question regarding Slope Sea reproduction is whether these mature individuals represent the putative Western stock or transient Mediterranean migrants. High-density single nucleotide polymorphism (SNP) markers confirm that Slope Sea larval cohorts exhibit genomic profiles more aligned with those of the Gulf of Mexico [Fraile et al. 2026]. With the Slope Sea situated in a highly dynamic oceanographic setting, recent findings identify complexities in connectivity and population structure that are not captured by the current two-stock management model [Rodríguez-Ezpeleta et al. 2019; Brody et al. 2020; Díaz-Arce et al. 2024; Fraile et al. 2026].

To evaluate the plausibility that Eastern-origin ABFT comprises the majority of fish that we sampled, we applied a binomial probability model for females (n = 42). If the Slope Sea spawning grounds were occupied predominantly by Eastern migrants, under a hypothetical 90% mixing rate where Western stock fish are rare anomalies, the probability of randomly sampling 42 females and finding zero Western-origin individuals is highly improbable (P < 0.01). Under a symmetric 50/50 mixed-stock hypothesis, the probability of capturing this specific contingent without encountering a single Western fish becomes infinitesimal (P ∼ 2.22 x 10^-13^). These modeled results demonstrate that classifying the sampled collection as an Eastern-dominated contingent is mathematically untenable.

Further validation of our full collection (combined sexes) is currently underway through Close-Kin Mark-Recapture analysis within the ICCAT GBYP framework [Fraile et al. 2026]. Regardless of the final assignment outcomes, the biological importance of the Slope Sea remains clear. Confirmation of a dominant Western origin would indicate a stock that is significantly more resilient and utilizes a broader geographic range for reproduction than currently recognized. Conversely, a higher-than-expected Eastern or mixed origin would reveal a level of trans-Atlantic connectivity and behavioral plasticity that fisheries management has failed to capture. Ultimately, resolving this genomic identity is the right step toward quantifying the current reproductive contribution of the Slope Sea to the Atlantic- wide population.

### Equivalence in Reproductive Contribution

The evidence of Slope Sea spawning calls for fundamental re-examination of the role smaller, sexually mature bluefin play in the stock. The “late-maturing” paradigm implicitly relies on the BOFFFF (“Big Old Fat Fertile Female Fish”) hypothesis, the premise that older, larger females produce disproportionately higher quality or quantity of progeny via maternal effects [Berkeley et al. 2004; Hixon et al. 2014]. However, comparative histological studies suggest that for ABFT, absolute fecundity is a function of allometric scaling rather than an age-dependent physiological advantage [Medina et al. 2002; Knapp et al. 2014; Corriero et al. 2020].

Crucially, relative batch fecundity remains consistent across size classes once physiological maturity is reached [Farley and Ohshimo 2019; Corriero et al. 2020]. We maintain that reproductive output in ABFT is primarily driven by individual energy allocation [Chapman et al. 2011] and the proximity to suitable oceanographic features, e.g., fronts and mesoscale eddies, that facilitate rapid caloric gain and optimal larval survival [Reglero et al. 2018; 2025]. The co-occurrence of mid-sized (123.3-190 cm SFL) and larger adults (Fig. 1) across the limited geographic extent of longline sets suggests that bluefin spawning contingents operate as mixed-age assemblages, subject to similar selective pressures. Consequently, current evidence does not support the assumption that larger individuals contribute disproportionately higher reproductive value per unit of spawning biomass than their mid-sized counterparts.

Schooling behavior of ABFT has been explored at observational levels [Lutcavage and Kraus 1995] that cannot determine intrinsic ethology. However, the migratory and reproductive knowledge attributed to older spawners likely functions as a shared social resource within a population [Couzin et al. 2005; Petit and Bon 2010]. As obligate schooling animals, where hydrodynamic constraints enforce strict size-sorting within individual schools [Pitcher 1986; Krause and Ruxton 2002], bluefin benefit from the collective intelligence of the group, particularly as cohorts coalesce into large, mixed-age foraging assemblages on common feeding grounds [Lutcavage and Kraus 1995; Fromentin and Powers 2005]. This is a critical but under-studied aspect of their life history. Through group consensus and social transmission, younger spawners can navigate to optimal spawning grounds and achieve reproductive success by leveraging established migratory pathways and collective learned memory.

### Reconciling Paradigms for Management

Our results demonstrate that the long-standing discrepancy between Western and Eastern stocks’ maturation schedules was likely an artifact of sampling constraints and a lack of understanding regarding ABFT physiological foundations. Maturity was assigned based on size-at-catch within an assumed, singular spawning ground, rather than direct biological analysis and physiological reality. By establishing an empirically derived L_50_ of 123.3 cm SFL, we establish a unified life-history model for the species, one that corrects past assumptions of sexual maturity, and transcends rigid management boundaries.

The presence of actively spawning mid-sized individuals in the Slope Sea supports a size-structured migration pattern that optimizes lifetime reproductive success across heterogeneous environments and energetic constraints [Chapman et al. 2011]. This demographic resilience, where diverse age cohorts utilize distinct oceanographic habitats, suggests *T. thynnus* is buffered against climate-driven shifts in larval survivorship.

Crucially, our estimated increase in spawning stock biomass (SSB) must not be conflated with surplus harvest capacity. Rather, these results identify a structural spawning contingent that was previously omitted from past and current assessment frameworks. As outlined in Box 1, the long-term sustainability of ABFT hinges on bridging the gap between large-scale statistical simulation and hard-won, empirical ground-truth.

The long-standing socio-economic impacts of these inaccurate portrayals of stock status and sustainability for Western Atlantic fleets and industry must also be considered. Size-at-maturity assumptions likely influenced selection of U.S. commercial and recreational fishery target size and category quotas, yielding severe economic consequences. For example, by 1993, the minimum regulatory size of targeted catch for US purse seine category was 206 cm CFL, and catches of smaller fish resulted in penalties and regulatory discards. In the mid 2000’s, the absence of large fish coupled with an influx of smaller fish into the Gulf of Maine led to zero purse seine catches, loss of crews, financial hardship, and, subsequently, the cessation of fishing as a livelihood for those license holders. Retrospective economic modeling of the structural impacts on U.S. and Canadian fleets under these regulatory regimes is highly warranted. Transitioning toward spatially explicit assessment models that reflect this underlying demographic diversity must now be an immediate assessment and management priority.

## Box 1. The Logistical Reality and Data Asymmetry of Bluefin Tuna Science

**Plate 1.**
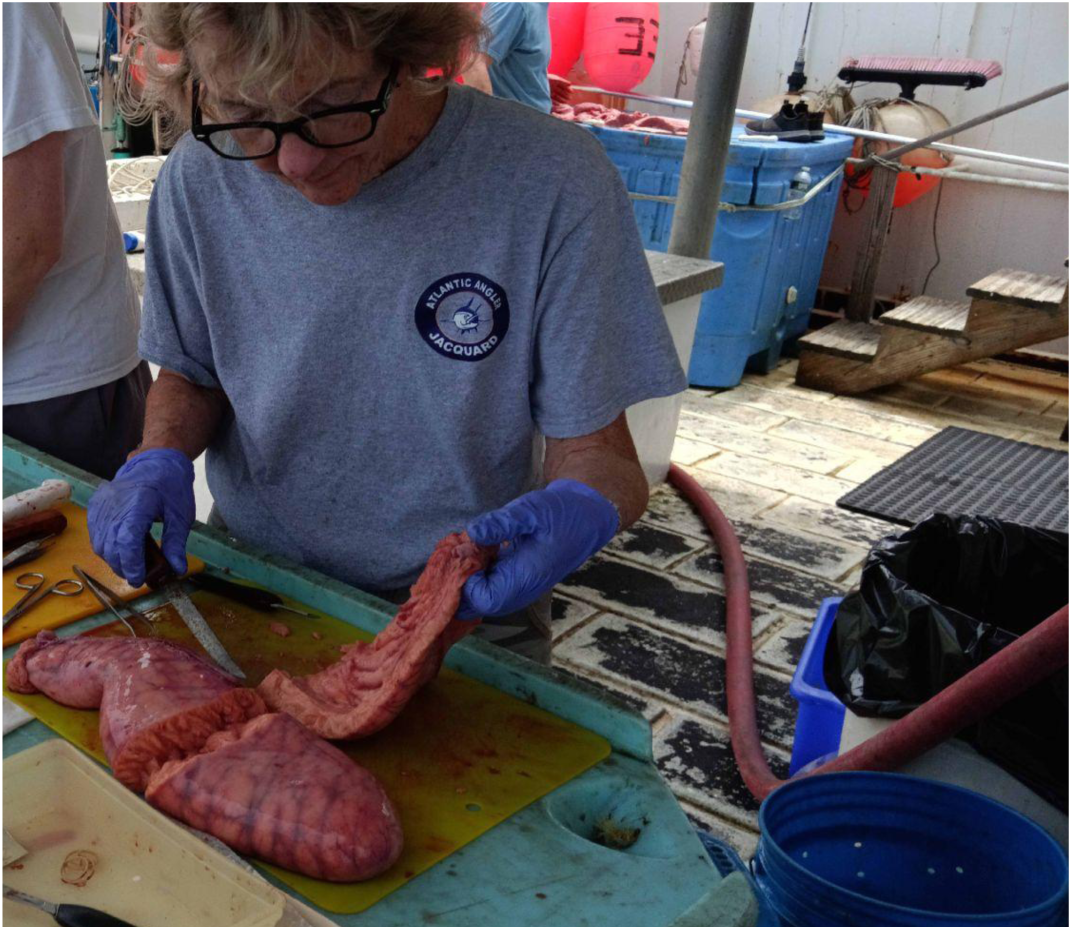
Aboard F/V *Eagle Eye II*, principal investigator and lead author, Molly Lutcavage (right) processed bluefin tuna gonad samples used in this study. Photo credit: Simon Gulak.

Histological analysis is the undisputed benchmark for tuna reproductive science, yet the path to obtaining these data is paved with practical challenges. The immense logistical, financial, and regulatory hurdles of field sampling do not fit nicely into the standardized frameworks of fisheries management; nonetheless, these realities dictate the availability and spatial resolution of the empirical data that underpin any informed management decision.

- **The Physiological Window:** Intercepting a spawning event requires precise timing, as histological markers like post-ovulatory follicles (POFs) degrade within 48 hours. Fortunately, maturity status is easier to assign. Because the transition to maturity is physiologically irreversible, our ability to define the maturity schedule remains robust even when the fleeting “smoking gun” of active spawning is missed.
- **The Environmental Gamble:** Field conditions dictate fish availability, yet both are notoriously unpredictable. During our June 2025 cruise, offshore temperatures were initially 16°C, forcing a full-day search, nearly 10% of our charter, just to locate fishable thermal fronts. This risk is inherent to expedition-based research. Our last attempt, dating back to 2001 in the Central North Atlantic, failed because it lacked the contemporary Slope Sea larval data and biotelemetry tracks of adults needed to time the spawning window. These “null results” are the invisible casualties of field biology, rarely persisting in our collective scientific memory, yet they explain precisely why histological ground-truth in the West Atlantic remains so sparse.
- **The Asymmetry of Indirect Inferences:** Because field sampling is so logistically fraught, research has gravitated toward “indirect proxies”, ranging from satellite telemetry to historical catch records. These offer vast, accessible datasets for “after-the-fact” statistical mining. While numerically attractive, the sheer volume of this indirect evidence often creates a “statistical gravity” that overshadows sparse, though more definitive biological observations, i.e., the gonads, or endocrine profiles that track the gonadal development. This imbalance is further entrenched by a reliance on simulation frameworks built upon initial assumptions that are rarely re-evaluated against new empirical, biological evidence. The result is that statistical power is unfortunately mistaken for biological accuracy.
- **Economic and Temporal Constraints:** Directed research cruises are governed by high fuel costs, labor intensity, and fixed duration. Ocean expeditions are rarely economical. Within this framework, time is the most restrictive commodity. Our modeling suggests that over 300 histological female samples are required to robustly define a Western maturity ogive (with traditional histology), a target virtually impossible to hit even with additional rounds of sampling. This financial and logistical barrier ensures that high-resolution ground-truth remains a rare luxury in fisheries science, where the “cost of biological accuracy” is often deemed too high to pay.
- **Regulatory Compliance:** In the US, scientific sampling for ABFT operates under strict enforcement and legal requirements where mandatory priorities are quota compliance and gear restrictions rather than experimental design. In this study, navigating these hurdles required a pragmatic collaboration between authorities and the community: by donating sampled carcasses to local food banks, we fulfilled permit requirements while ensuring the resource was utilized rather than wasted as mandated regulatory discards. This represents a real-world complexity that no statistical framework can model or appreciate.

Ultimately, these factors converge into a formidable “sampling filter.” The scarcity of histological data confirming spawning fish in the pelagic Western Atlantic is not necessarily indicative of their lack, but rather the immense difficulty of being in the right place, at the right time, with the right people and gear. The “late-maturing” Western paradigm was less a product of biological observation than a direct consequence of the logistical path of least resistance. This Box serves as a reminder that while statistical and simulation frameworks provide an indispensable tool for fisheries management, they are only as reliable as the sparse, hard-won biological “ground-truths” that provide the necessary reality checks.

## Declaration of Generative AI and AI-assisted Technologies in the Writing Process

During the preparation of this manuscript, the authors utilized the Gemini 3 Flash (Web interface) to assist in the computational modeling. Specifically, the AI was employed to implement the Python-based simulation code for the spawning stock biomass sensitivity and cumulative distribution analysis.

Following the use of these tools, the authors reviewed and edited the generated content. The authors take full responsibility for the accuracy of the biological assumptions, the integrity of the data analysis, and the final conclusions presented in the manuscript. To ensure reproducibility, the Python code used for the simulation is included in the Supplementary Materials.

## Reproducibility Script (Python)

See annotated code in https://docs.google.com/document/d/1Nd_8mzgIrPfnSHYbagSC2×8c7JwaVnGbmf_Y1ayBJDg

